# Analyses of symbiotic bacterial communities in the plant pest *Bemisia tabaci* reveal high prevalence of *Candidatus* Hemipteriphilus asiaticus on the African continent

**DOI:** 10.1101/2021.10.06.463217

**Authors:** Laurence Mouton, Hélène Henri, Rahim Romba, Zainab Belgaidi, Olivier Gnankiné, Fabrice Vavre

## Abstract

Microbial symbionts are widespread in insects and some of them have been associated to adaptive changes. Primary symbionts (P-symbionts) have a nutritional role that allows their hosts to feed on unbalanced diets (plant sap, wood, blood). Most of them have undergone genome reduction, but their genomes still retain genes involved in pathways that are necessary to synthesize the nutrients that their hosts need. However, in some P-symbionts, essential pathways are incomplete and secondary symbionts (S-symbionts) are required to complete parts of their degenerated functions. The P-symbiont of the phloem sap-feeder *Bemisia tabaci, Candidatus* Portiera aleyrodidarium, lacks genes involved in the synthesis of vitamins, cofactors, and also of some essential amino-acids. Seven S-symbionts have been detected in the *B*. tabaci species complex. Phenotypic and genomic analyses have revealed various effects, from reproductive manipulation to fitness benefits, notably some of them have complementary metabolic capabilities to *Candidatus* Portiera aleyrodidarium, suggesting that their presence may be obligatory. In order to get the full picture of the symbiotic community of this pest, we investigated, through metabarcoding approaches, the symbiont content of individuals from Burkina Faso, a West African country where *B. tabaci* induces severe crop damage. While no new putative *B. tabaci* S-symbiont was identified, *Candidatus* Hemipteriphilus asiaticus, a symbiont only described in *B. tabaci* populations from Asia, was detected for the first time on this continent. Phylogenetic analyses however reveal that it is a different strain than the reference found in Asia. Specific diagnostic PCRs showed a high prevalence of these S-symbionts and especially of *Candidatus* Hemipteriphilus asiaticus in different genetic groups. These results suggest that *Candidatus* Hemipteriphilus asiaticus may affect the biology of *B. tabaci* and provide fitness advantage in some *B. tabaci* populations.

## Introduction

Many insects feed on nutrient-imbalanced diet (plant sap, wood, blood), and there is strong experimental evidence that microbial symbionts can promote utilization of these resources by the synthesis of essential nutrients, like amino-acids and vitamins (for review see 1). In hemipterans, these obligatory symbionts, also called primary symbionts (P-symbionts), are intracellular, strictly maternally inherited and, for most of them, have evolved with their hosts for millions of years (review in 2). They are often housed inside specialized cells, bacteriocytes, within a dedicated organ, the bacteriome, localized in the host’s abdomen, which constitutes a stable environment for the symbionts and facilitates their transmission to offspring (3, 4). As a consequence of this lifestyle, their genomes are extremely reduced (5, 6). Their minimal genomes retain genes involved in pathways that complement essential nutrients lacking in their host’ diets (7). However, in some bacterial symbionts, gene repertoire seems insufficient to meet the metabolic demand of their hosts (review in 8). For example, in aphids, in the subfamily Lachninae, the P-Symbiont *Buchnera aphidicola* has lost its ability to synthesize tryptophan and riboflavin (9).

These deficiencies can be compensated by the acquisition of alternative symbionts that can mediate equivalent functions. Indeed, in addition to their P-symbiont, insects often harbour secondary symbionts (S-symbionts). These symbionts are predominantly vertically transmitted but can also be horizontally transmitted, and have a wide range of effects from mutualism to parasitism (review in 10). Several studies have shown that these co-resident S-symbionts can complement the metabolic network of the P-symbionts leading to an inter-dependency between the symbiotic partners. Thus, all the members of the aphid subfamily Lachninae depend on a second co-obligate symbiont to complement specific gene losses of the P-symbiont *Buchnera* (9, 11; review in 12). These co-symbionts are numerous with eleven identified up to now: ten γ-proteobacteria and one α-proteobacteria. This inter-dependency between the symbiotic partners is not restricted to aphids and *Buchnera*, it has been described in other hemiptera like in Cicadas where the P-symbiont *Sulcia* has almost always been detected with one or more co-obligate symbionts (12).

Like other phloem-sap feeders, the whitefly *Bemisia tabaci* harbours a P-symbiont, *Candidatus* Portiera aleyrodidarum (13), that synthetizes essential nutrients. However, *Ca*. Portiera aleyrodidarum has a tiny genome, around 355kb (14-17) and, while this γ-proteobacterium has the capacity to synthesize carotenoids and most essential amino-acids (14), it lacks almost all the genes involved in the synthesis of vitamins and cofactors. Moreover, pathways involved in the synthesis of some essential amino acids are incomplete (18). Bacteria belonging to seven genera of S-symbionts have been identified in the cryptic species complex of *B. tabaci*: *Hamiltonella, Arsenophonus, Cardinium, Rickettsia, Wolbachia, Fritschea* and *Hemipteriphilus* (19-22). They are all localized in the same bacteriocytes as the P-symbiont but some can also be found in the hemolymph (23). Their roles remain poorly understood but range from reproductive parasitism (24) to fitness benefits such as thermal tolerance (25, 26). Moreover, the analysis of their genomes suggests that some of them could play a nutritional role. For example, *Hamiltonella* can provide vitamins and cofactors, and could also complete the missing steps of the lysine pathway of *Ca*. Portiera aleyrodidarum (18). Because of these complementations, presence of S-symbionts is expected in all whitefly individuals. In a sampling performed in West Africa in 2007 and 2009, *B. tabaci* individuals were indeed predominantly found infected with S-symbionts, but in some populations no S-symbiont were recorded (27). In this former study, prevalence of S-symbionts was determined with an *a priori* method, *i. e*. through PCRs using specific primers targeting the six symbionts identified in *B. tabaci* at the time, thus leaving the possibility that other, undescribed endosymbionts were present. Since, a seventh S-symbiont, *Candidatus* Hemipteriphilus asiaticus (hereafter “*Ca*. Hemipteriphilus asiaticus”), has been described in *B. tabaci* (19-20). This makes it possible that *Ca*. Hemipteriphilus asiaticus, as well as other bacterial symbionts, are in fact present in the S-symbiont free *B. tabaci* individuals.

In order to get a full picture of the symbiont diversity, we investigated the symbiont content of individuals sampled in Burkina Faso (West Africa) using a metabarcoding approach. In this country, *B. tabaci* is a pest of primary importance, with a severe impact on economic activity (28). Indeed, *B. tabaci* is a cryptic species complex (42 species reported till now based on a 657bp portion of the mitochondrial cytochrome oxidase 1 (*mtCOI*) DNA sequence: 29-33), and in Burkina Faso individuals belong either to SSA (Sub-Saharan Africa), ASL (Africa Silver-Leafing) or MED (Meditteranean) species (34). Using universal bacterial primer sets we detected few bacterial species and, more importantly, no new putative S-symbiont. However, the S-symbionts known in *B. tabaci* were found, except *Fritschea*, and notably *Ca*. Hemipteriphilus asiaticus that is described for the first time in Africa. Interestingly, phylogenetic analyses revealed that this *Ca*. Hemipteriphilus asiaticus differs from the reference strain identified in Asia (20). In addition, diagnostic PCRs revealed high prevalence of S-symbionts in these populations, and notably *Ca*. Hemipteriphilus asiaticus in ASL and MED individuals, which questions its possible role in the biology of *B. tabaci*.

## Results

Among the 630 individuals sampled in Burkina Faso in 2015 and 2016, the majority, almost 84%, belonged to the MED species, more especially to the MED-Q1 genetic group (81% ; 3% belonged to MED-Q3). ASL and SSA (more precisely SSA2) genetic groups represented 10.5% and 3% of individuals respectively (34).

### Bacterial community characterization

The bacterial community of *B. tabaci* collected in Burkina Faso was characterized on 72 individuals by a metabarcoding approach, without *a priori* assumptions, using universal bacterial primers targeting the 16S rRNA gene and an Illumina sequencing technology. Between 107,000 to 927,000 reads were obtained per sample (average: 215,000 reads ± 9,420). The majority of the reads belonged to the known *B. tabaci* P- and S-symbionts and, in most field individuals, sequences from *Ca*. Portiera aleyrodidarum constituted the majority of reads (up to 98.1%; 71.2% on average ± 2.9%; see “table_level6” in Dryad, https://doi.org/10.5061/dryad.547d7wm91, and Figure 1). We detected bacterial taxa belonging to the genera *Acinectobacter* in few samples at low abundances since they represented between 0 and 7,4% of the sequences obtained per individual, both in field and lab samples. They may represent gut bacteria or contaminations. Overall, no new S-symbiont has been detected.

**Figure 1.**
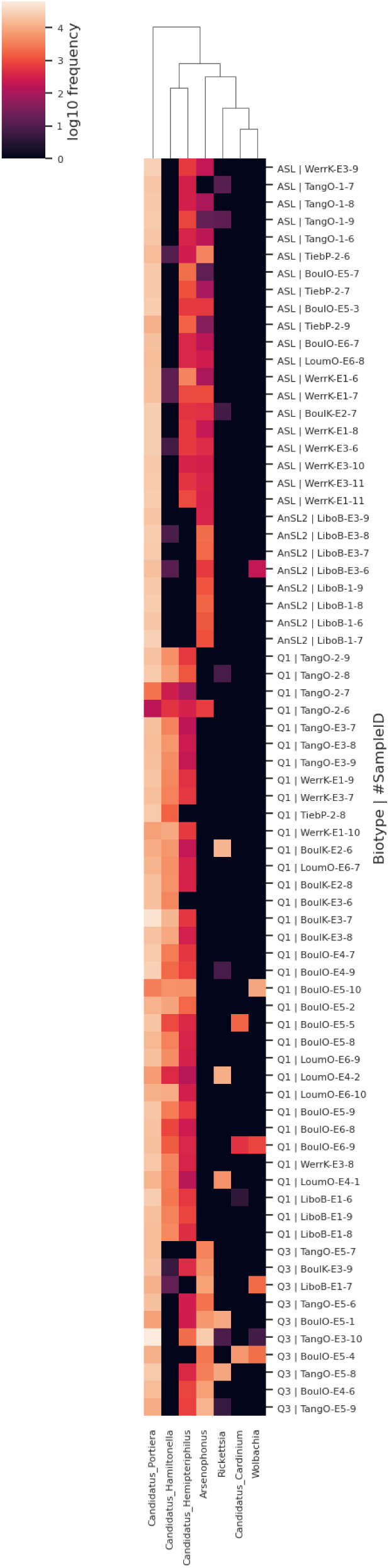
Heat-map showing the major taxa identified in the metabarcoding analysis. Heatmap showing differences in bacterial communities based on taxonomic classifications of DNA 16S amplicons generated in QIIME 2 using the SSU SILVA 138 taxonomy. The heatmap was generated from the log-transformed relative abundance values of the 7 major taxa at the genus level (level-6). The relative abundance of each genus is indicated by a gradient of color from darkest (low abundance) to lightest color (high abundance).

All the known *B. tabaci* S-symbionts have been found except *Fritschea*. However, it is not surprising since this S-symbiont seems to be scarce worldwide in *B. tabaci* (35). Moreover, in a previous survey done in West Africa using specific primers *Fritschea* was not found (27). As already shown, there is a link between symbiotic bacterial communities and genetic groups: *Hamiltonella* was found to be the most common S-symbiont in MED-Q1, and *Arsenophonus* in MED-Q3 and SSA2 individuals. *Ca*. Hemipteriphilus asiaticus is the only S-symbiont presents in all genetic groups except SSA2 (*i*.*e*. in MED-Q1, MED-Q3 and ASL). It was not detected in individuals of the laboratory lines.

### Phylogenetic analysis of *Ca*. Hemipteriphilus asiaticus

Twenty individuals positive for *Ca*. Hemipteriphilus asiaticus, belonging to MED-Q1, MED-Q3 and ASL genetic groups, were used for phylogenetic analysis. They originated from eight localities and, when possible, in each locality, the three genetic groups were represented (Table 1).

**Table 1.** Samples used for the phylogenetic analyses of *Ca*. Hemipteriphilus asiaticus.

| Locality | Host plant | Sample name | Biotype |
| --- | --- | --- | --- |
| Tanghin/Ouagadougou | Eggplant | TangO22 | MED-Q1 |
|  | *Lantana camara* | TangOE56 | MED-Q3 |
|  | Potato | TangO14 | ASL |
|  | Tomato | TangOE35 | ASL |
| Tiebele/Pô | Chilli pepper | TiebPE11 | MED-Q1 |
|  | Potato | TiebP13 | ASL |
| Boulmiougou/Ouagadougou | Bell pepper | BoulOE33 | MED-Q1 |
|  | Cucumber | BoulOE44 | MED-Q3 |
|  | Cucumber | BoulOE46 | MED-Q3 |
|  | Zucchini | BoulOE51 | MED-Q3 |
|  | Tomato | BoulOE15 | ASL |
| Boulbi/Komsilga | Eggplant | BoulKE11 | MED-Q1 |
|  | Eggplant | BoulKE33 | MED-Q3 |
|  | Zucchini | BoulKE39 | MED-Q3 |
| Koubri/Kombissiri | Bell pepper | KoubKE11 | MED-Q1 |
|  | Zucchini | KoubK45 | MED-Q3 |
| Werra/Koudougou | Eggplant | WerrKE24 | MED-Q1 |
|  | Tomato | WerrE11 | ASL |
| Loumbila/Oubritenga | Chilli pepper | LoumOE14 | MED-Q1 |
| Bonyolo/Réo | Eggplant | BonyRE13 | MED-Q1 |

The 16S rRNA sequences (483bp) obtained with the new primers designed in the present study (Table 2) were 100% identical in all the individuals (*i. e*. 8 MED-Q1, 7 MED-Q3 and 5 ASL). This sequence showed 100% similarity with the ones of *Ca*. Hemipteriphilus asiaticus endosymbiont and *Rickettsia* of *B. tabaci* available in the genbank database (Blast done in December 2021). 16S rRNA gene is highly conserved (36) we thus designed new primers on two other loci, *GltA* and *GroEL*, using the sequences of the *Ca*. Hemipteriphilus asiaticus isolate YH-ZHJ available in Genbank (20). The 190bp sequences of *GltA* were identical in all our 20 samples, but one substitution was found in the *GroEL* sequences (269bp) between MED individuals (Q1 and Q3) and ASL individuals, whatever their sampling locality.

**Table 2.**
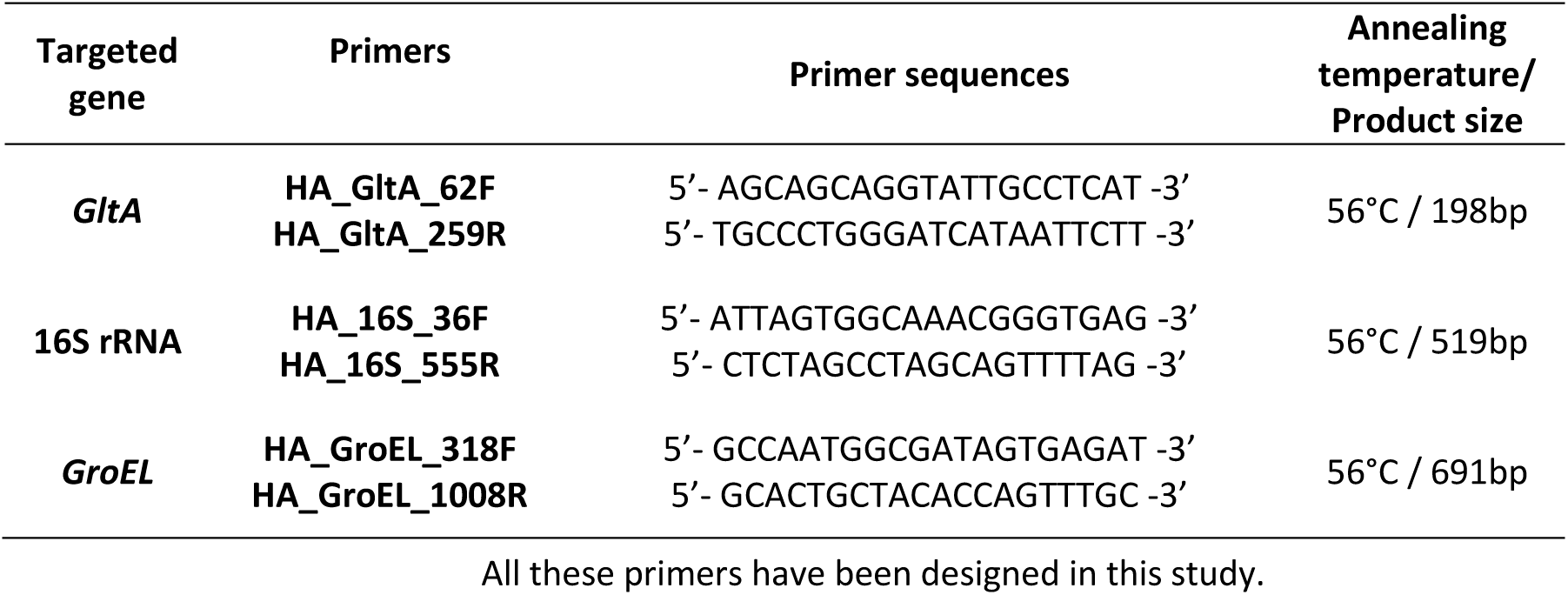
PCR primers and conditions used for phylogenetic analysis of *Ca*. Hemipteriphilus asiaticus.

Analyses of the concatenated sequences obtained for the three loci (942bp in total) revealed ∼97% identity between the *Ca*. Hemipteriphilus asiaticus strains identified in Burkina Faso and the YH-ZHJ reference isolate from the China *B. tabaci* species (between 1 to 13 different bases according to the gene). Topologies of the trees for *GroEL* and *GltA* were similar to the one of the concatenated tree (*16s rRNA* is not informative since the 483bp sequence of *Ca*. Hemipteriphilus asiaticus is 100% similar to *Rickettsia*).

We also analysed the phylogenetic relationships of *Ca*. Hemipteriphilus asiaticus and other close Rickettsiales, *Rickettsia* and *Sitobion miscanthi L Type Symbiont* SMLS. The trees constructed with maximum-likelihood and Bayesian inference methods are identical and showed that *Ca*. Hemipteriphilus asiaticus is closer to SMLS than to *Rickettsia* (Figure 2), which is concordant with analyses of Bing *et al*. (20) and Li *et al*. (37). This phylogenetic analysis thus confirms the presence of *Ca*. Hemipteriphilus asiaticus in MED-Q1, MED-Q3 and ASL genetic groups in Burkina Faso. It also reveals that they represent different strains than the one found in Asia, and that the strain found in ASL differs slightly from the strains in MED-Q1 and MED-Q3.

**Figure 2.**
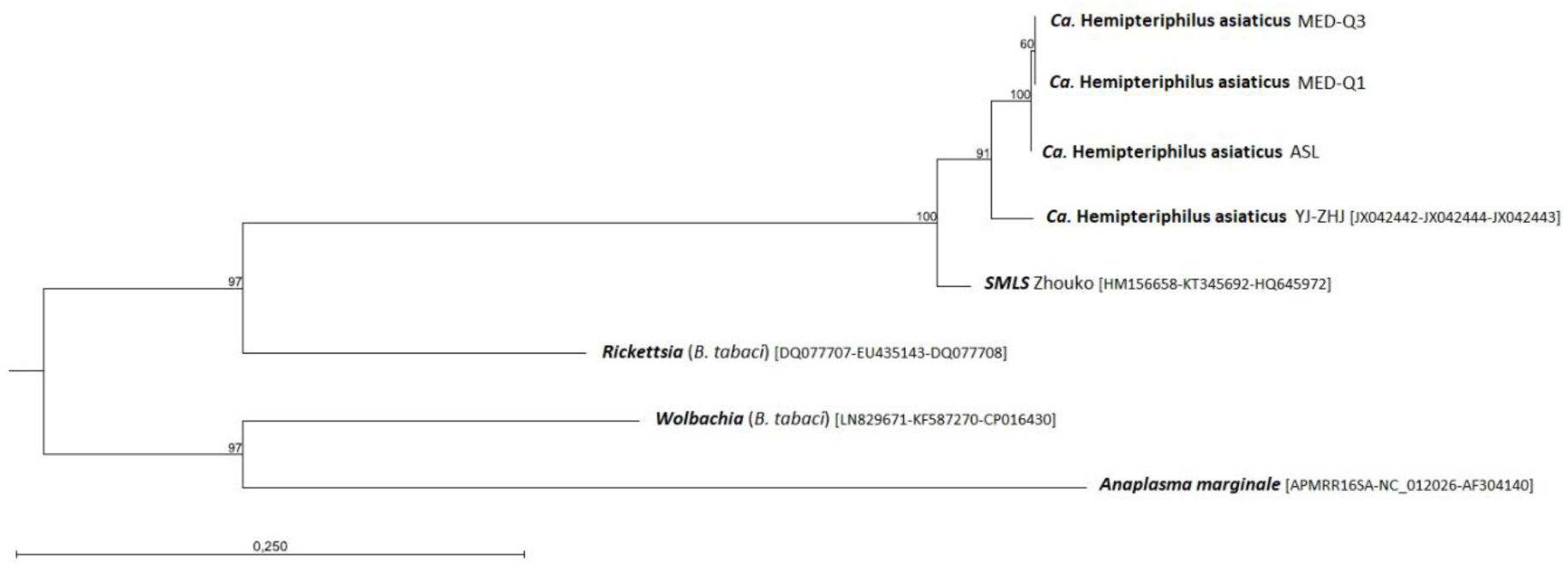
Phylogeny constructed using Maximum Likelihood analysis (model GTR + G + T) based on the trimmed and concatenated sequences of three genes, 16S rRNA (483bp), *GroEL* (269bp) and *GltA* (190bp) (942bp in total). Boostrap values are shown at the nodes (100 replicates). Sequences obtained from *Ca*. Hemipteripilus asiaticus found in the present study in *B. tabaci* from Burkina Faso (indicated by the biotype of their host, MED-Q1, MED-Q3, ASL) were compared to the YH-ZHJ strain found in China1 biotype individuals of *B. tabaci* from China and to other closely related symbionts belonging to the Rickettsiales family, *Rickettsia* and *Sitobion miscanthi L Type Symbiont* (SMLS). The genbank accession numbers of the sequences obtained from other studies are indicated in the brackets.

### Distribution and prevalence of bacterial endosymbionts in field populations

The presence of the P- and S-symbionts was checked in 334 individuals from nine localities in Burkina Faso (Figure 3) and several host plants (vegetables, ornamental plants and weeds) by specific diagnostic qPCRs.

**Figure 3.**
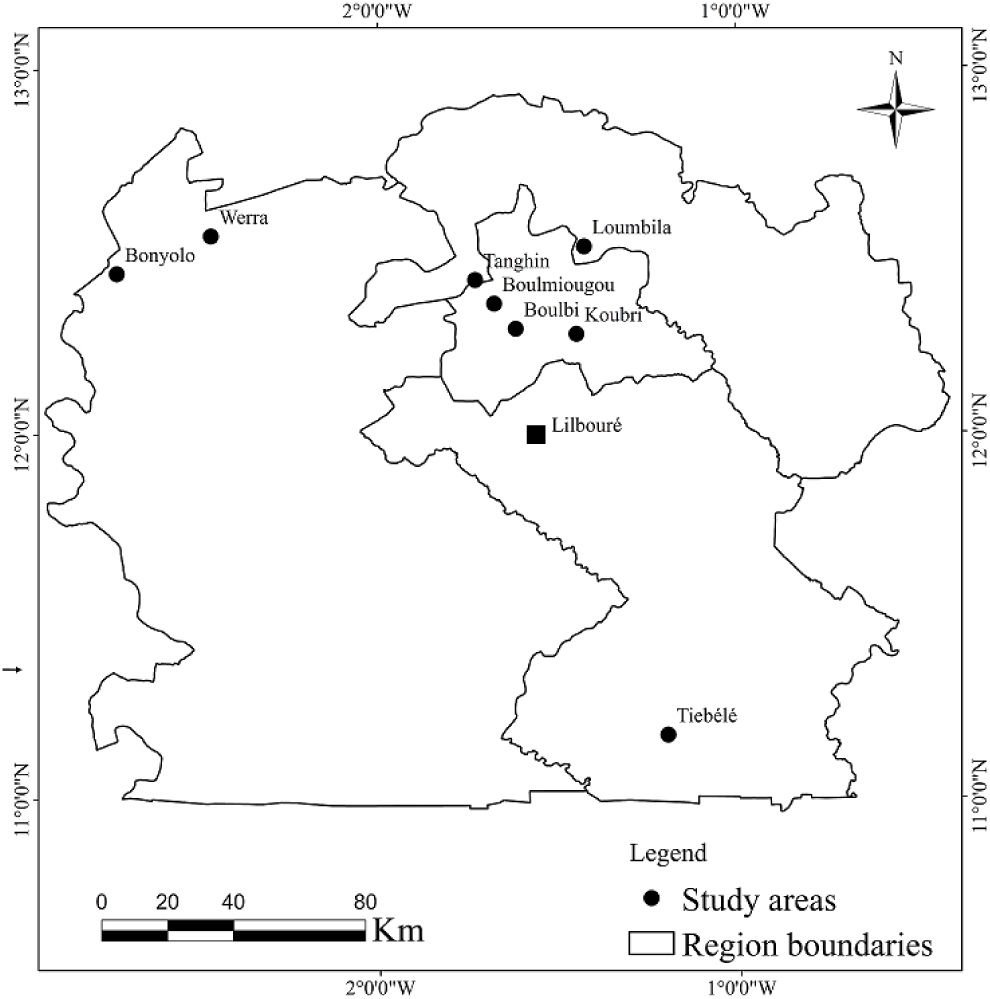
Sampling localities in Burkina Faso. Both prevalence of symbionts and phylogenetic analysis of *Ca*. Hemipteriphilus asiaticus were done on individuals from the nine localities indicated except Lilbouré (indicated by a square) for which no sample was used for the phylogenies.

All the S-symbionts described so far in *B. tabaci* were targeted except *Fritschea bemisiae* because this bacterium was not found in the 16S rRNA metabarcoding analysis. The infection status of individuals for the P-symbiont as well as the S-symbionts (*Hamiltonella, Arsenophonus, Cardinium, Rickettsia, Wolbachia, Ca*. Hemipteriphilus asiaticus) are presented in Figure 4. *Ca*. Portiera aleyrodidarum was found in all but five individuals from different sampling sites. PCRs done on the actine host gene as well as the detection of S-symbionts ensured the extraction quality, but we cannot exclude that the quantity of *Ca*. Portiera aleyrodidarum was under the real-time PCR detection threshold in these samples. The fact that these five individuals all belong to MED-Q1 can be explained by the high prevalence of this genetic group (262/334).

**Figure 4.**
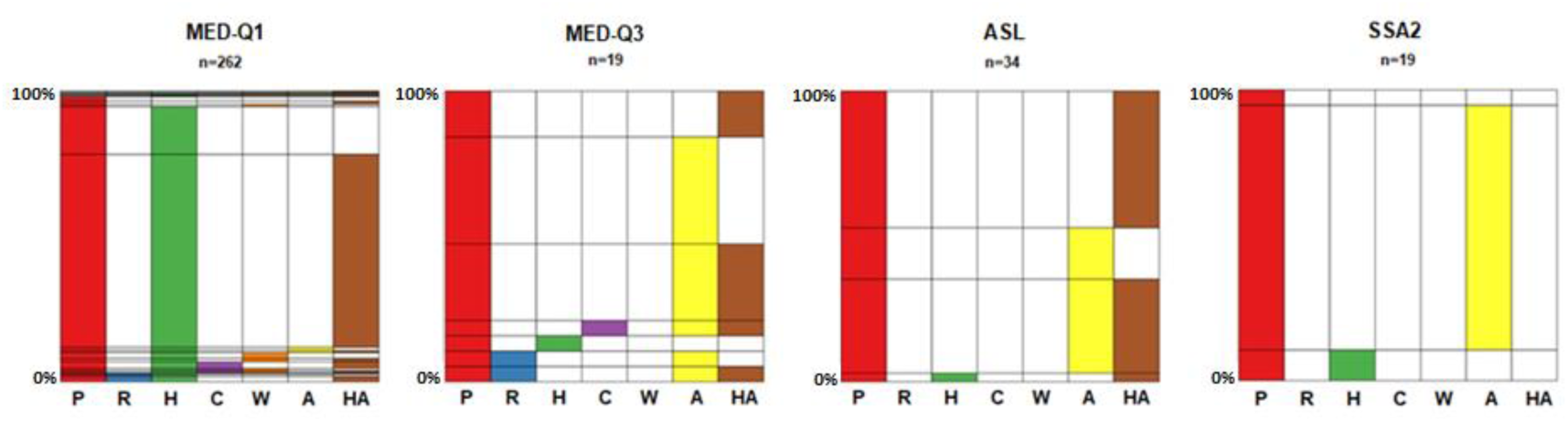
Infection status of *Bemisia tabaci* individuals collected in Burkina Faso according to their biotype, determined through specific qPCRs. Each graph corresponds to one biotype, with the different bacterial symbionts shown on the x-axis. Each bacterium is represented by a colour (red: *Ca*. Portiera aleyrodidarum (P), blue: *Rickettsia* (R), green: *Hamiltonella* (H), purple: *Cardinium* (C), orange: *Wolbachia* (W), yellow: *Arsenophonus* (A), brown: *Ca*. Hemipteriphilus asiaticus (HA)) and the corresponding coloured bar indicates its prevalence. On the y-axis host individuals are ranked and grouped together according to their infection status: when the graph is read horizontally, the colour combinations represent individuals sharing the same symbiotic community. n indicates the number of individuals checked.

More than 98% of individuals harboured at least one S-symbiont and, as expected, the prevalence of the S-symbionts genera depended on the genetic group (Fisher’s Exact Test, P = 0.0005; see data analysis of 21). *Arsenophonus* is the most frequent symbiont in SSA2 and MED-Q3 individuals (89% and 79% respectively), while 96% of individuals belonging to MED-Q1 harbour *Hamiltonella*. Interestingly, *Ca*. Hemipteriphilus asiaticus is dominant in ASL samples with 82% of individuals infected, while 50% harbour *Arsenophonus*. More generally *Ca*. Hemipteriphilus asiaticus is very frequent in all biotypes, except SSA2 in which it has not been found. For the first time, this symbiont is described in MED-Q1, MED-Q3 and ASL genetic groups. In all of them, its prevalence is high: 77% in MED-Q1, 53% in MED-Q3 and 82% in ASL. Globally, the symbiotic composition in these four genetic groups corresponds to what is known in literature (meta-analysis in Zchori-Fein *et al*. (21); see Gnankiné *et al*. (27) for previous data obtained in Burkina Faso), except for *Ca*. Hemipteriphilus asiaticus. The symbiotic composition is not influenced by the locality for MED-Q1 (Fisher’s Exact Test, P = 0.1414), which is the only genetic group found in all the localities (Figure S1).

Co-infection by several S-symbionts is frequent (69%), mostly double-infections which represent 90% of multiple infections. However, the presence of three or even four S-symbionts within the same host individual has also been detected (20 and 2 individuals respectively; Figure 4). An exception is the biotype SSA2 in which only single infections have been found. Some associations of symbionts are frequent and others never found. We never found *Rickettsia*-*Cardinium, Rickettsia*-*Wolbachia* and *Wolbachia*-*Arsenophonus* combinations. On the other hand, in the MED-Q1 individuals *Hamiltonella* and *Ca*. Hemipteriphilus asiaticus co-occur frequently: this assemblage represents 76% of bi-infections in this genetic group (193/253). In MED-Q3 and ASL, *Ca*. Hemipteriphilus asiaticus is often associated with *Arsenophonus* (58% and 92% respectively). These results show that *Ca*. Hemipteriphilus asiaticus often co-infect individuals with another S-symbiont, especially the two more frequent S-symbionts, that are also the ones confined to bacteriocytes.

### Influence of *Ca*. Hemipteriphilus asiaticus on the P-symbiont density

As it is the first time that *Ca*. Hemipteriphilus asiaticus has been detected in *B. tabaci* individuals from the African continent, we aimed at determining whether its presence has an impact on the P-symbiont. We thus compared the density of *Ca*. Portiera aleyrodidarum in presence and in absence of *Ca*. Hemipteriphilus asiaticus (Figure 5). Results indicated that the presence of *Ca*. Hemipteriphilus asiaticus does not affect the density of *Ca*. Portiera aleyrodidarum (n=282, Wilcox rank test, W=8364, P=0.0847).

**Figure 5.**
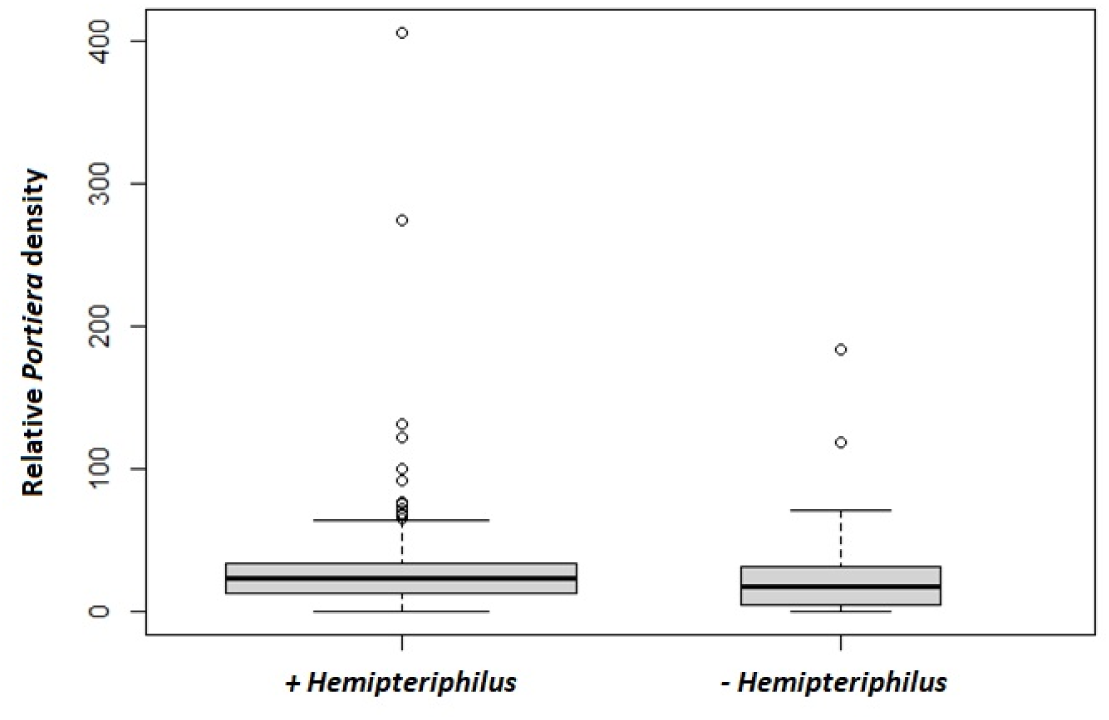
*Ca*. Portiera aleyrodidarum densities according to the presence of *Ca*. Hemipteriphilus asiaticus. Relative *Ca*. Portiera aleyrodidarum (*Portiera* in the figure) densities (ratio of the number of copies of *Ca*. Portiera aleyrodidarum (16S rRNA) and host (actine) genes) in absence (-; n=69) or presence (+ ; n=213) of *Ca*. Hemipteriphilus asiaticus (*Hemipteriphilus* in the figure) within the same host individuals. The width of the box plots reflects the number of samples for each modality.

## Discussion

The whitefly *B. tabaci* is one of the most devastating agricultural pests worldwide. In Africa, damage induced by this species complex is huge and results in severe impacts on the economic activity of many countries. Effective control requires understanding its ability to spread and determining factors involved in its important polyphagy. In this context, heritable bacterial symbionts are of primary importance since they may provide their hosts with important ecological traits. In the present study, we aimed at describing the symbiotic bacterial communities, diversity and prevalence in *B. tabaci* populations from Burkina Faso (West Africa). In this survey, several *B. tabaci* genetic groups were found: ASL (Africa Silver-Leafing) and MED (Meditteranean, MED-Q1 and MED-Q3), as previously reported by Gnankiné *et al*. (27), and, in addition, SSA (Sub-Saharan Africa), that was detected for the first time in this country, but only in one locality (Lilboure), and only on one host plant, cassava (34).

The metabarcoding analysis of the bacterial symbionts did not reveal the presence of symbionts not yet described in *B. tabaci*. However, *Ca*. Hemipteriphilus asiaticus is described for the first time in Africa: it was detected in all the genetic groups found in Burkina Faso except SSA, *i. e*. MED-Q1, MED-Q3 and ASL. *Ca*. Hemipteriphilus asiaticus has been described for the first time in 2013 by Bing *et al*. (20) in *B. tabaci* samples from China belonging to the China1 biotype. Since then, it has also been found in China2, Asia (I and II), Indian Ocean and SSA (SSA6) genetic groups, but only in countries of the Asian continent: Indian and Pakistan (22, 38, 39). In the present study, *Ca*. Hemipteriphilus asiaticus was detected in the African continent, with a very high prevalence in the MED and ASL genetic groups from Burkina Faso: 53% in MED-Q3, 77% in MED-Q1, reaching 82% in ASL. On the other hand, it has not been found in SSA2. Since its description, *Ca*. Hemipteriphilus asiaticus has been relatively under-studied. To our knowledge, only two field surveys have been done so far. They both found the presence of *Ca*. Hemipteriphilus asiaticus in Asia I and II species but with high differences in prevalence: the infection rate was high in Central India (89%: 51 out of 57 individuals checked were infected; 38), but lower in Pakistan (between 5% and 39%; 22). Based on all these results, the presence of *Ca*. Hemipteriphilus asiaticus should be sought in population studies when *a priori* methods based on specific PCRs are used to describe the bacterial community associated with the *B. tabaci* complex species. This is especially true as we describe here that *Ca*. Hemipteriphilus asiaticus can also infect the worldly distributed MED-Q1.

Our phylogenetic analyses on *Ca*. Hemipteriphilus asiaticus based on three genes, 16S rRNA, *GroEL* and *GltA* revealed that 3% of nucleotide sites differ between the MED strain identified in the present study and the reference YH-ZHJ isolate described in the China *B. tabaci* species (20). It also revealed a substitution in the *GroEL* sequence between the *Ca*. Hemipteriphilus asiaticus found in MED and in ASL, yet found in sympatry. Therefore, the present data reveal the existence of polymorphism in the *Ca*. Hemipteriphilus asiaticus genus with at least three strains present in *B. tabaci*. Naturally, in further studies, an extended sampling should be done in more continents, countries, on more species/genetic groups, and more molecular markers should be developed in order to get a more accurate idea of the diversity of *Ca*. Hemipteriphilus asiaticus strains in this complex species. Anyway, *Ca*. Hemipteriphilus asiaticus is not an isolated case: several S-symbionts (*Wolbachia, Arsenophonus, Cardinium* and *Rickettsia)* are represented by more than one strain in *B. tabaci* (35, 40), with up to 6 phylogenetic groups identified for *Arsenophonus* (41).

To date the influence of *Ca*. Hemipteriphilus asiaticus on its host is not known. In *B. tabaci*, it has been suggested that some S-symbionts could play a nutritional role, in collaboration with the P-symbiont. Indeed, several data suggest that *Hamiltonella* and *Arsenophonus* could have an impact on the *B. tabaci* metabolism and dietary requirements. These two S-symbionts are almost fixed in some genetic groups (27, 41, present study). For instance, *Hamiltonella* is widespread in MEAM1 and MED-Q1 (review in 21 and 35). Moreover, *Hamiltonella* possesses some genes involved in amino-acid biosynthesis pathways that are lost or non-functional in the P-symbiont (18). In addition, recent experiments demonstrated that this S-symbiont can supply *B. tabaci* with the production of B vitamins (42-43). Even if there is no evidence of fixation in any genetic group in the field, *Ca*. Hemipteriphilus asiaticus could confer a benefit to its host under some environmental conditions, for example, according to the nutritional quality of the host plants. Indeed, previous research demonstrated that aphid performance is associated with the amino-acid composition of the phloem sap (44-45). It could explain why *Ca*. Hemipteriphilus asiaticus was not present in the SSA species which, contrary to MED and ASL which have been found in several host plant species, has only been detected on cassava in Burkina Faso (34). Analysis of the genome of *Ca*. Hemipteriphilus asiaticus, with a special focus on genes involved in metabolic pathways, would give further insight into its putative nutritional role.

The presence of another closely relative S-symbiont, *Sitobion miscanthi L Type Symbiont* (SMLS), has also been recently highlighted in the aphid *Sitobion miscanthi* (46). It also belongs to the Rickettsiaceae family and, similarly to *Ca*. Hemipteriphilus asiaticus, is widely distributed in some populations of its host (see survey in China in 47). It has been suggested that SMLS could stimulate the proliferation of the P-symbiont *Buchnera* and thus improve the aphids’ fitness. Indeed, *Buchnera*’s density is significantly higher in SMLS-infected individuals and laboratory experiments revealed that infected individuals show higher values of some fitness traits (37). However, our results did not reveal any influence of *Ca*. Hemipteriphilus asiaticus on the *Ca*. Portiera aleyrodidarum density. Clearly, life history traits should be measured on *Ca*. Hemipteriphilus asiaticus-infected and -free *B. tabaci* individuals. Anyway, the high frequency of these two newly reported S-symbionts in some field populations of these phloemophagous insects could suggest they bring benefit to their hosts. Further research on the infection dynamics of these S-symbionts, and their role on their host phenotype and adaptation are needed.

Compared to the previous field survey done in Burkina Faso by Gnankiné *et al*. (27), data on the infection with S-symbionts are highly similar except for the presence of *Ca*. Hemipteriphilus asiaticus, which was not described at that time. In particular, the infection rate involving at least one S-symbiont were higher than 90% in the two studies. In MED-Q1 genetic group, *Hamiltonella* is the more frequent S-symbiont (96% in the present study, 89% in 27) while *Arsenophonus* is the most common bacteria in MED-Q3 and ASL individuals (previous study/present study, respectively 93%/79% and 40%/50%). Interestingly, *Hamiltonella* and *Arsenophonus* are mutually exclusive which is not the case of *Ca*. Hemipteriphilus asiaticus that is often found in co-infection with these S-symbionts: it was involved in almost 80% of co-infection by two S-symbionts.

## Conclusion

In summary, these data confirmed the variability of the symbiotic community in the *B. tabaci* complex species despite its high temporal stability in populations from Burkina Faso, and reveal the presence of another player whose role deserves to be studied. The stability and the high indicence of S-symbionts in *B. tabaci*, together with genomic studies, suggest that they can have central roles in shaping the fitness of this pest in different environments. Understanding their possible contribution to successfull invasion, widespread distribution, and more generally, on the population dynamics of this whitefly is critical for the implementation of effective pest management programs.

## Methods

### Sampling

Sampling was done in nine localities in Burkina Faso, Western Africa (Figure 3), in March and April (dry season) 2015 and 2016 on vegetables, ornamental plants and weeds as described in Romba *et al*. (34). The species and biotypes (hereafter genetic groups) of the individuals (adults) were determined in Romba *et al*. (34) according to the PCR-RFLP method developed in Henri *et al*. (48). Three species were found, ASL (Africa Silver-Leafing), SSA (Sub-Saharan Africa) and MED (Mediterranean). Within the MED species, two genetic groups, MED-Q1 and MED-Q3, were identified.

### Characterization of the bacterial community

Seventy-two field-collected whiteflies from Burkina Faso were used to characterize the bacterial community. They were chosen in order that all genetic groups, all localities and all host plants were represented (Table 1). We also included 18 adults coming from laboratory lines belonging to MED-Q1 and MED-Q2 genetic groups reared for years in the « Laboratoire de Biométrie et Biologie Evolutive ». The universal bacterial primer set 341F-*TCG-TCG-GCA-GCG-TCA-GAT-GTG-TAT-AAG-AGA-CAG***-**CCT-ACG-GGN-GGC-WGC-AG and 805R-*GTC-TCG-TGG-GCT-CGG-AGA-TGT-GTA-TAA-GAG-ACA-G*GA-CTA-CHV-GGG-TAT-CTA-ATC-C was used to amplify 464 bp of the V3-V4 hypervariable regions of the 16S rRNA gene (49). The primers were synthesized with overhang adapters (in italic) for index attachment and Illumina sequencing adapters. For each sample, which consisted in one individual, triplicates were performed, consisting of three PCR reactions using the KAPA ReadyMix (KAPA) containing 200nM of each primer, 12.5µL of KAPA Hifi HotStart Ready Mix and 2.5 µL of DNA template, in a final volume of 25 µL. The conditions of reactions were 95°C for 3min followed by 25 cycles of 95°C for 30sec, 55°C for 30sec and 72°C for 30sec. Then a final elongation was done at 72°C for 5min. Each amplification product was checked on an agarose gel to verify that there was specific amplification only. For some of them, a bioanalyzer verification was also performed. Then, all the replicates were pooled by sample before purification and to proceed to the further preparation of the library according to the protocol outlined by Illumina (« 16S metagenomic Sequencing Library Preparation »), n°15044223 Rev.B. The pooled library was PE-sequenced using the Illumina MiSeq reagent kit version 3 for 600 cycles (2×300pb) by Biofidal (Vaulx enVelin, France).

The sequencing data in FASTQ format were processed and analyzed with the QIIME2 software suite version 2021.11 (50). The raw Illumina reads were imported into QIIME2, demultiplexed, and then denoised, trimmed and filtered with DADA2 pipeline to remove noisy and chimeric sequences, to construct denoised paired-end sequences and to dereplicate them (51). This produced a table containing representative sequences also called amplicon sequence variant or ASV. The taxonomy assignment was then performed by using feature-classifier classify-sklearn.. For that, reads from the Silva 138 SSURef NR99 reference database were extracted to match on the primer set 341F/805R. The Naive Bayes classifier has been trained before the classification (52-53). A taxa barplot has been done on all the data with “qiime taxa bar plot” (except chloroplast sequences that have been removed; Dryad, https://doi.org/10.5061/dryad.547d7wm91) and a heatmap has been produced only on field samples for major taxa with “qiime feature table heatmap” (Figure 1).

### Primer design

We designed new primers for *Ca*. Hemipteriphilus asiaticus using sequences available in Genbank (20) with the Primer3 software (https://bioinfo.ut.ee/primer3-0.4.0/). We aligned sequences of *Ca*. Hemipteriphilus asiaticus with sequences of *Rickettsia* from *B. tabaci* in order to design primers specific to *Ca*. Hemipteriphilus asiaticus that do not amplify sequences from *Rickettsia*. These primers were tested on individuals harbouring *Rickettsia* but not *Ca*. Hemipteriphilus asiaticus and we didn’t detect any amplification. Multiple sequence alignments were done using the MUSCLE algorithm (54) implemented in CLC DNA Workbench 8.0 (CLC Bio). Three sets of primers targeting three different genes of *Ca*. Hemipteriphilus asiaticus, 16S rRNA, the citrate synthase *GltA* and the chaperonin *GroEL* were designed (Table 2).

### Phylogenetic analysis of *Ca*. Hemipteriphilus asiaticus strains

PCRs targeting *GltA, GroEL* and 16S rRNA genes of *Ca*. Hemipteriphilus asiaticus were carried out on 20 positive samples (Table 1) with sets of primers designed specifically for this study (Table 2). DNA was amplified in a final volume of 25μL containing 200μM dNTPs, 200nM of each primer, 0.5IU DreamTaq DNA polymerase (ThermoFisher) and 2μL of DNA. All PCR amplifications were performed under the following conditions: initial denaturation at 95°C for 2min followed by 35 cycles at 94°C for 30sec, 56°C for 30sec, 72°C for 1min and a final extension at 72°C for 10min. PCR products were sequenced using the Sanger method by the platform Biofidal (Vaulx en Velin, France).

The nucleotide polymorphism was analyzed by aligning the sequences obtained using the MUSCLE algorithm (54) implemented in CLC DNA Workbench 8.0 (CLC Bio) and inspecting them by eye. Moreover, we studied the phylogenetic relationships of the *Ca*. Hemipteriphilus asiaticus strains found in our samples with the reference strain described in *B. tabaci* (isolate YH-ZHJ; 20) as well as other symbionts belonging to the Rickettsiales family, *Rickettsia, Orientia tsutsumagushi* and *Sitobion miscanthi L Type Symbiont* (SMLS). Phylogenetic trees were constructed with CLC DNA Workbench 8.0 (CLC Bio) using maximum-likelihood method for each sequence separately and for the concatenated data set (substitution model: GTR + G + T, choosed by using the “Model Testing” tool of CLC DNA Workbench 8.0). The robustness of nodes was assessed with 100 bootstrap replicates. We also constructed a phylogenetic tree with Bayesian inference using the program MrBayes (version 3.2.6) and a GTR+G model (55). For these concatenated gene dataset, 20000 generations were run and the first 25% of these were discarded as burn-in.

### Prevalence of bacterial endosymbionts

Three hundred and thirty four individuals were screened for the presence of the P-symbiont *Ca*. Portiera aleyrodidarum and six secondary symbionts, *Hamiltonella defensa, Arsenophonus* sp., *Cardinium hertigii, Rickettsia* sp., *Wolbachia pipientis* and *Ca*. Hemipteriphilus asiaticus (hereafter *Portiera, Hamiltonella, Arsenophonus, Cardinium, Rickettsia, Wolbachia* and *Ca*. Hemipteriphilus asiaticus) using specific PCR primers (Table 3). We did not check for the presence of *Fritschea bemisiae* in these individuals because this bacterium was not detected in the 16S rRNA metabarcoding analysis. We also amplified one host gene (actin) using the following primers: wf-B actin-For:5’-TCT-TCC-AGC-CAT-CCT-TCT-TG-3’ and wf-B actin-Rev: 5-CGG-TGA-TTT-CCT-TCT-GCA-TT-3’ to ensure the quality of DNA extractions. DNA was amplified in a final volume of 10μL containing 5μL of Sso Advanced SYBR Green Supermix (Bio-Rad), 2μL of water, 0.5μL of each primer (final concentration of 500nM) and 2μL of DNA samples. The reaction conditions for amplification were 95°C for 30sec followed by 40 cycles of 95°C for 10sec, 55°C to 63°C (according to the primers’ set) for 30sec (see Table 3 ; 63°C for Actine) and 72°C for 30sec. The specificity of the amplified products was controlled by checking melting curves (65°C to 95°C). To assess the efficiency of the reaction, standard curves were plotted using dilutions of previously amplified and purified PCR products. Amplification and detection of DNA were done on the real-time CFX96 instrument (Bio-Rad).

**Table 3.**
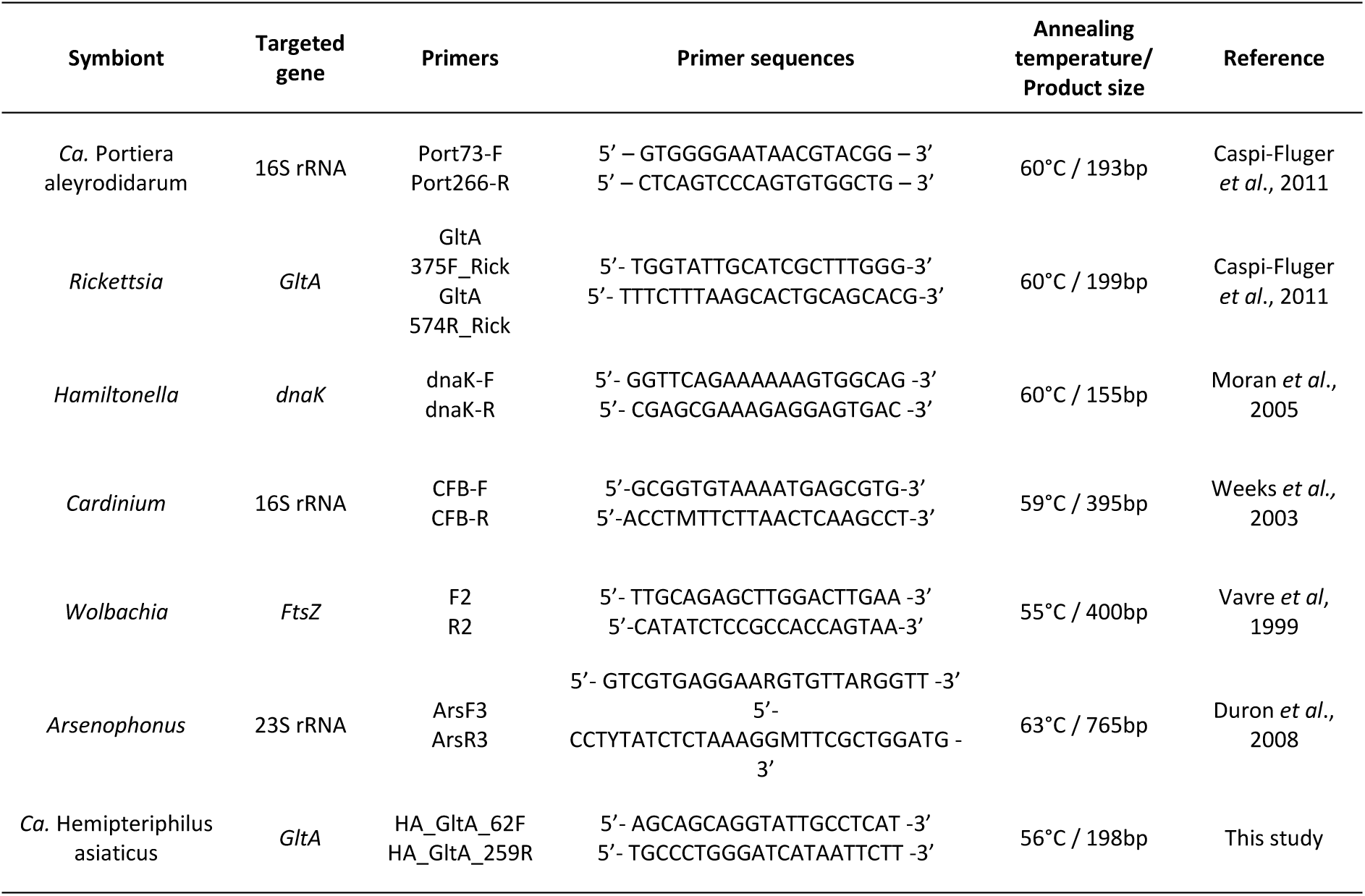
PCR primers and conditions used for symbionts’ screening (qPCR)

### Statistical analysis

Data analysis were performed using the R statistical software version 3.2.2 (http://www.R-project.org). The infection status of *B. tabaci* individuals was graphically represented using the Mondrian shiny application (Siberchicot, Charif, Terraz &Vavre: https://cran.r-project.org/web/packages/Mondrian/).

## Data accessibility

Nucleotide sequences obtained in this study are accessible in GenBank database under accession numbers MW353022 and MW353023 for *GroEL* (MED and ASL species respectively), MW343733 for 16S rRNA and MW353021 for *GltA*.

All datasets generated and analyzed on the bacterial community characterization are available in Dryad at: https://doi.org/10.5061/dryad.547d7wm91.

## Acknowledgements

This study was funded by Campus France, the Agence Nationale de la Recherche (HMicMac, ANR16-CE02-00014) and the Scientific Breakthrough Project Micro-be-have (Microbial impact on insect behavior) of Université de Lyon, within the programme ‘Investissements d’Avenir’ (ANR-11-IDEX-0007; ANR-16-IDEX-0005). The authors thank Agnès Nguyen from the Biofidal company for her precious advice and collaboration on the metabarcoding part. Version 3 of this preprint has been peer-reviewed and recommended by Peer Community In Zoology (https://doi.org/10.24072/pci.zool.100011).

## Supplementary material

**Figure S1.**
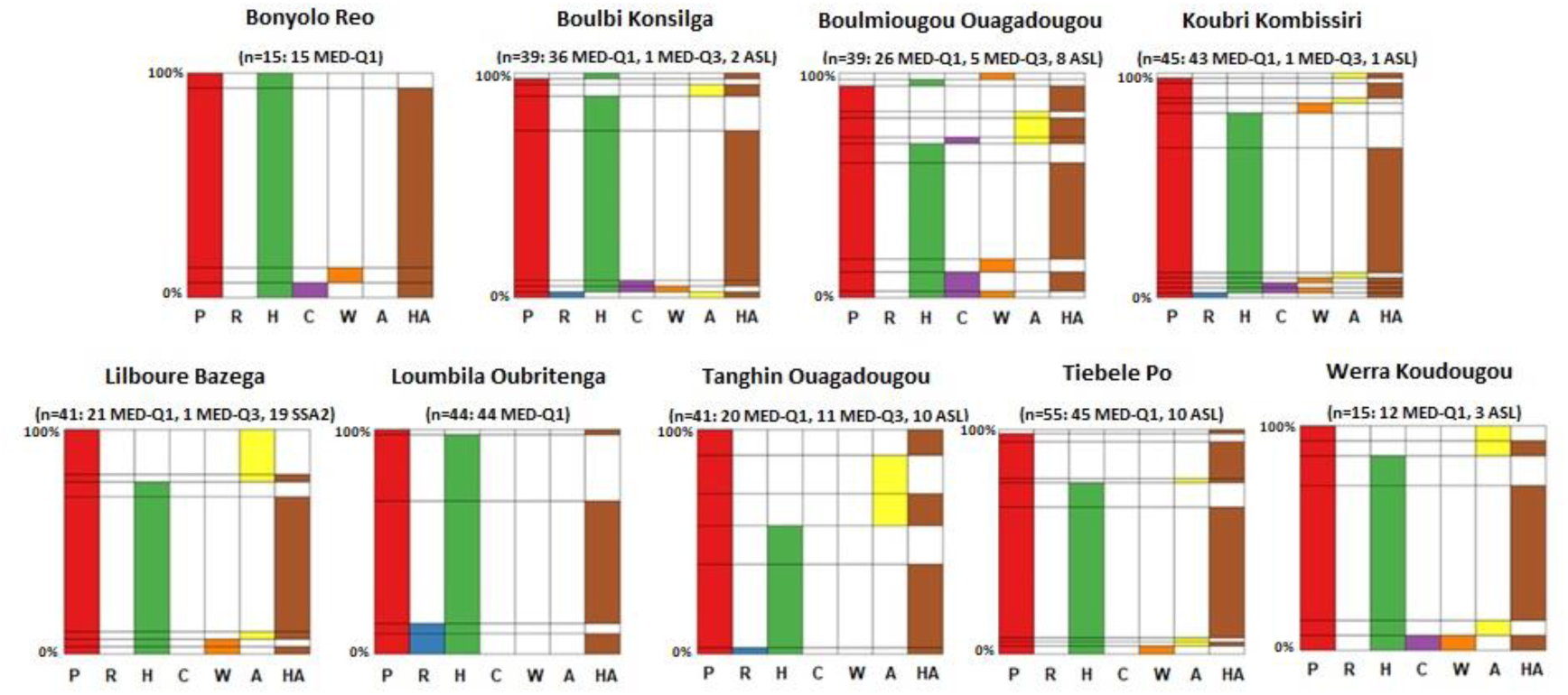
Infection status of *Bemisia tabaci* individuals collected in Burkina Faso according to the locality, determined through specific qPCRs. Each graph corresponds to one locality in Burkina Faso, with the different bacterial symbionts shown on the x-axis. Each bacterium is represented by a colour (red: *Ca*. Portiera aleyrodidarum (P), blue: *Rickettsia* (R), green: *Hamiltonella* (H), purple: *Cardinium* (C), orange: *Wolbachia* (W), yellow: *Arsenophonus* (A), brown: *Ca*. Hemipteriphilus asiaticus (HA)) and the corresponding coloured bar indicates its prevalence. On the y-axis host individuals are ranked and grouped together according to their infection status: when the graph is read horizontally, the colour combinations represent individuals sharing the same symbiotic community. n indicates the number of individuals checked. The number of individuals of each biotype is also indicated.

## References

1. Douglas AE. 2009. The microbial dimension in insect nutritional ecology. Funct Ecol 23: 38–47. https://doi.org/10.1111/j.1365-2435.2008.01442.x

2. Moran NA, McCutcheon JP, Nakabachi A. 2008. Genomics and evolution of heritable bacterial symbionts. Annu Rev Genet 42: 165–90. https://doi.org/10.1146/annurev.genet.41.110306.130119

3. Buchner P. 1965. Endosymbioses of animals with plant microorganisms. New York : Interspecience Publisher.

4. Baumann, P. 2005. Biology of Endosymbionts of Plant Sap-Sucking Insects. Ann Rev Microbiol 59: 155–89. https://doi.org/10.1146/annurev.micro.59.030804.121041

5. McCutcheon JP, Moran NA. 2011. Extreme genome reduction in symbiotic bacteria. Nat Rev Microbiol. 10: 13–26. https://doi.org/10.1038/nrmicro2670

6. Moran NA, Bennett GM. 2014. The tiniest tiny genomes. Annu Rev Microbiol 68: 195–215. https://doi.org/10.1146/annurev-micro-091213-112901

7. Gil R, Silva FJ, Pereto J, Moya A. 2004. Determination of the core of a minimal bacterial gene set. Microbiol Mol Biol R 68: 518–537. https://doi.org/10.1128/MMBR.68.3.518-537.2004

8. Latorre A, Manzano-Marín A. 2017. Dissecting genome reduction and trait loss in insect endosymbionts. Ann NY Acad Sci 1389: 52–75. https://doi.org/10.1111/nyas.13222

9. Manzano-Marín A, Simon JC, Latorre A. 2016. Reinventing the wheel and making it round again: evolutionary convergence in Buchnera-Serratia symbiotic consortia between the distantly related Lachninae Aphids Tuberolachnus salignus and Cinara cedri. Genome Biol Evol 8: 1440–1458. https://doi.org/10.1093/gbe/evw085

10. Ferrari J, Vavre F. 2011. Bacterial symbionts in insects or the story of communities affecting communities. Philos Trans R Soc Lond B Biol Sci 366: 1389–1400. https://doi.org/10.1098/rstb.2010.0226

11. Meseguer AS, Manzano-Marín A, Coeur d’Acier A, Clamens AL, Godefroid M, Jousselin E. 2017. Buchnera has changed flatmate but the repeated replacement of co-obligate symbionts is not associated with the ecological expansions of their aphid hosts. Mol Ecol 26: 2363–2378. https://doi.org/10.1111/mec.13910

12. Gil R, Latorre A. 2019. Unity makes strength: a review on mutualistic symbiosis in representative insect clades. Life 9: 21. https://doi.org/10.3390/life9010021

13. Thao MLL, Baumann P. 2004. Evolutionary relationships of primary prokaryotic endosymbionts of whiteflies and their hosts. App Env Microbiol 70: 3401–3406. https://doi.org/10.1128/AEM.70.6.3401-3406.2004

14. Santos-Garcia D, Farnier PA, Beitia F, Zchori-Fein E, Vavre F, Mouton L, Moya A, Latorre A, Silva FJ. 2012. Complete genome sequence of “Candidatus Portiera aleyrodidarum” BT-QVLC, an obligate symbiont that supplies amino acids and carotenoids to Bemisia tabaci. J Bacteriol 194: 6654–6655. https://doi.org/10.1128/JB.01793-12

15. Sloan DB, Moran NA. 2012. Genome reduction and co-evolution between the primary and secondary bacterial symbionts of psyllids. Mol Biol Evol 29: 3781–3792. https://doi.org/10.1093/molbev/mss180

16. Sloan DB, Moran NA. 2013. The evolution of genomic instability in the obligate endosymbionts of whiteflies. Genome Biol Evol 5: 783–793. https://doi.org/10.1093/gbe/evt044

17. Jiang ZF, Xia F, Johnson KW, Brown CD, Bartom E, Tuteja JH, Stevens R, Grossman RL, Brumin M, White KP, Ghanim M. 2013. Comparison of the genome sequences of “Candidatus Portiera aleyrodidarum” primary endosymbionts of the whitefly Bemisia tabaci B and Q Biotypes. Appl Env Microbiol 79: 1757–1759. https://doi.org/10.1128/AEM.02976-12

18. Rao Q, Rollat-Farnier PA, Zhu DT, Santos-Garcia D, Silva FJ, Moya A, Latorre A, Klein C, Vavre F, Sagot MF, Liu SS, Wang XW, Mouton L. 2015. Genome reduction and metabolic complementation of the dual endosymbionts in the whitefly Bemisia tabaci. BMC Genomics 16: 226. https://doi.org/10.1186/s12864-015-1379-6

19. Bing XL, Ruan YM, Rao Q, Wang XW, Liu SS. 2012. Diversity of secondary endosymbionts among different putative species of the whitefly Bemisia tabaci. Insect Sci 20: 194–206. https://doi.org/10.1111/j.1744-7917.2012.01522.x

20. Bing XL, Yang J, Zchori-Fein E, Wang XW, Liu SS. 2013. Characterization of a newly discovered symbiont of the whitefly Bemisia tabaci (Hemiptera : Aleyrodidae). Appl Env Microbiol 79: 569–575. https://doi.org/10.1128/AEM.03030-12

21. Zchori-Fein E, Lahav T, Freilich S. 2014. Variations in the identity and complexity of endosymbiont combinations in whitefly hosts. Front Microbiol 5: 310. https://doi.org/10.3389/fmicb.2014.00310

22. Wang HL, Lei T, Xia WQ, Cameron SL, Liu YQ, Zhang Z, Gowda MMN, Navas-Castillo J, Omongo CA, Delatte H, Lee KY, Patel MV, Krause-Sakate R, Ng J, Wu SL, Fiallo-Olivé E, Liu SS, Colvin J, Wang XW. 2019. Insight into the microbial world of Bemisia tabaci cryptic species complex and its relationships with its host. Sci Rep 9: 6568. https://doi.org/10.1038/s41598-019-42793-8

23. Gottlieb Y, Ghanim M, Gueguen G, Kontsedalov S, Vavre F, Fleury F, Zchori-Fein E. 2008. Inherited intracellular ecosystem: symbiotic bacteria share bacteriocytes in whiteflies. FASEB J. 22: 2591–2599. https://doi.org/10.1096/fj.07-101162

24. Lv N, Peng J, Chen XY, Guo CF, Sang W, Wang XM, Ahmed MZ, Xu YY, Qiu BL. 2020. Antagonistic interaction between male-killing and cytoplasmic incompatibility induced by Cardinium and Wolbachia in the whitefly, Bemisia tabaci. Insect Sci 28: 330–346. https://doi.org/10.1111/1744-7917.12793

25. Brumin M, Kontsedalov S, Ghanim M. 2011. Rickettsia influences thermotolerance in the whitefly Bemisia tabaci B biotype. Insect Sci 18: 57–66. https://doi.org/10.1111/j.1744-7917.2010.01396.x

26. Yang K, Yuan MY, Liu Y, Guo CL, Liu TX, Zhang YJ, Chu D. 2021. First evidence for thermal tolerance benefits of the bacterial symbiont Cardinium in an invasive whitefly, Bemisia tabaci. Pest Manag Sci 77: 5021–5031. https://doi.org/10.1002/ps.6543

27. Gnankiné O, Mouton L, Henri H, Terraz G, Houndeté T, Martin T, Vavre F, Fleury F. 2013. Distribution of Bemisia tabaci biotypes (Homoptera: Aleyrodidae) and their associated symbiotic bacteria on host plants in western Africa. Ins Cons Div 6: 411–421. https://doi.org/10.1111/j.1752-4598.2012.00206.x

28. Gnankiné O, Traore D, Sanon A, Traore NS, Ouédraogo AP. 2007. Traitements insecticides et dynamique des populations de Bemisia tabaci en culture cotonnière au Burkina Faso. Cahiers Agriculture 16: 101–109. https://doi.org/10.1684/agr.2007.0081

29. Dinsdale A, Cook L, Riginos C, Buckley YM, De Barro P. 2010. Refined global analysis of Bemisia tabaci (Hemiptera: Sternorrhynca: Aleyrodoidae: Aleyrodidae) mitochondrial cytochrome oxidase 1 to identify species level genomic boundaries. Ann Entomol Soc Am 103: 196–208. https://doi.org/10.1603/AN09061

30. De Barro PJ, Liu SS, Boykin LM, Dinsdale AB. 2011. Bemisia tabaci: a statement of species status. Ann Rev Entomol 56: 1–19. https://doi.org/10.1146/annurev-ento-112408-085504

31. Firdaus S, Vosman B, Hidayati N, Supena EDJ, Visser RGF, van Heusden AW. 2013. The Bemisia tabaci species complex : additions from different parts of the world. Insect Sci 20: 723–733. https://doi.org/10.1111/1744-7917.12001

32. Roopa HK, Asokan R, Rebijith KB, Hande RH, Mahmood R, Kumar NK. 2015. Prevalence of a new genetic group, MEAM-K, of the whitefly Bemisia tabaci (Hemiptera : Aleyrodidae) in Karnataka, India, as evident from mtCOI sequences. Florida Entomol 98: 1062–1071. https://doi.org/10.1653/024.098.0409

33. Hu J, Zhang X, Jiang Z, Zhang F, Liu Y, Liu Z, Zhang Z. 2018. New putative cryptic species detection and genetic network analysis of Bemisia tabaci (Hemiptera: Aleyrodidae) in China based on mitochondrial COI sequences. Mitochondrial DNA Part A. 29: 474–484. https://doi.org/10.1080/24701394.2017.1307974

34. Romba R, Gnankine O, Drabo SF, Tiendrebeogo F, Henri H, Mouton L, Vavre F. 2018. Abundance of Bemisia tabaci Gennadius (Hemiptera : Aleyrodidae) and its parasitoids on vegetables and cassava plants in Burkina Faso (West Africa). Ecol Evol 8: 6091–6103. https://doi.org/10.1002/ece3.4078

35. Kanakala S, Ghanim M. 2019. Global genetic diversity and geographical distribution of Bemisia tabaci and its bacterial endosymbionts. PLoS One 14: e0213946. https://doi.org/10.1371/journal.pone.0213946

36. Barth Reller L, Weinstein MP, Petti CA. 2007. Detection and identification of microorganisms by gene amplification and sequencing. Clin inf dis 44: 1108–14. https://doi.org/10.1086/512818

37. Li T, Wu XJ, Jiang YL, Zhang L, Duan Y, Miao J, Gong ZJ, Wu YQ. 2016. The genetic diversity of SMLS (Sitobion miscanthi L type symbiont) and its effect on the fitness, mitochondrial DNA diversity and Buchnera aphidicola dynamic of wheat aphid, Sitobion miscanthi (Hemiptera : Aphididae). Mol Ecol 25: 3142–3151. https://doi.org/10.1111/mec.13669

38. Ansari PG, Singh RK, Kaushik S, Krishna A, Wada T, Noda H. 2017. Detection of symbionts and virus in the whitefly Bemisia tabaci (Hemiptera: Aleyrodidae), vector of the Mungbean yellow mosaic India virus in Central India. Appl Entomol Zool 52: 567–579. https://doi.org/10.1007/s13355-017-0510-3

39. Paredes-Montero JR, Zia-Ur-Rehman M, Hameed U, Haider MS, Herrmann HW, Brown JK. 2020. Genetic variability, community structure, and horizontal transfer of endosymbionts among three Asia II-Bemisia tabaci mitotypes in Pakistan. Ecol Evol 10: 2928–2943. https://doi.org/10.1002/ece3.6107

40. Gueguen G, Vavre F, Gnankine O, Peterschmitt M, Charif D, Chiel E, Gottlieb Y, Ghanim M, Zchori-Fein E, Fleury F. 2010. Endosymbiont metacommunities, mtDNA diversity and the evolution of the Bemisia tabaci (Hemiptera: Aleyrodidae) species complex. Mol Ecol 19: 4365–4376. https://doi.org/10.1111/j.1365-294X.2010.04775.x

41. Mouton L, Thierry M, Henri H, Baudin R, Gnankiné O, Reynaud B, Zchori-Fein E, Becker N, Fleury F, Delatte H. 2012. MLST diversity and recombination evidence in Arsenophonus symbionts of the Bemisia tabaci species complex. BMC Microbiol 12(Suppl. 1): S10. https://doi.org/10.1186/1471-2180-12-S1-S10

42. Ren FR, Sun X, Wang TY, Yao YL, Huang YZ, Zhang X, Luan JB. 2020. Biotin provisioning by horizontally transferred genes from bacteria confers animal fitness benefits. ISME J. 14: 2542–2553. https://doi.org/10.1038/s41396-020-0704-5

43. Wang YB, Ren FR, Yao YL, Sun X, Walling LL, Li NN, Bai B, Bao XY, Xu XR, Luan JB. 2020. Intracellular symbionts drive sex ratio in the whitefly by facilitating fertilization and provisioning of B vitamins. ISME 14: 2923–2935. https://doi.org/10.1038/s41396-020-0717-0

44. Sandström J, Pettersson J. 1994. Amino acid composition of phloem sap and the relation to intraspecific variation in pea aphid (Acyrthosiphon pisum) performance. J Insect Phys 40: 947–955.

45. Stout MJ, Thaler JS, Thomma BPHJ. 2006. Plant-mediated interactions between pathogenic microorganisms and herbivorous arthropods. Ann Rev Entomol 51: 663–689. https://doi.org/10.1146/annurev.ento.51.110104.151117

46. Li T, Xiao JH, Xu ZH, Murphy RW, Huang DW. 2011. A possibly new Rickettsia-like genus symbiont is found in Chinese wheat pest aphid, Sitobion miscanthi (Hemiptera : Aphididae). J Inv Pathol 106: 418–421. https://doi.org/10.1016/j.jip.2010.12.003

47. Li T, Xiao JH, Wu YQ, Huang DW. 2014. Diversity of bacterial symbionts in populations of Sitobion miscanthi (Hemiptera : Aphididae) in China. Env Entomol 43: 605–611. https://doi.org/10.1603/EN13229

48. Henri H, Terraz G, Gnankiné O, Fleury F, Mouton L. 2013. Molecular characterization of genetic diversity within the Africa/Middle East/Asia Minor and Sub-Saharan Africa groups of the Bemisia tabaci species complex. Int J Pest Management 59: 329–338. https://doi.org/10.1080/09670874.2013.869374

49. Klindworth A, Pruesse E, Schweer T, Peplies J, Quast C, Horn M, Glöckner FO. 2013. Evaluation of general 16S ribosomal RNA gene PCR primers for classical and next-generation sequencing-based diversity studies. Nucleic Acids Res. 41: e1. https://doi.org/10.1093/nar/gks808

50. Bolyen E, Rideout JR, Dillon MR, Bokulich NA, Abnet CC, Al-Ghalith GA, et al. 2019. Reproducible, interactive, scalable and extensible microbiome data science using QIIME 2. Nat Biotechnol 37:852–857. https://doi.org/10.1038/s41587-019-0209-9

51. Callahan BJ, McMurdie PJ, Rosen MJ, Han AW, Johnson AJA, Holmes SP. 2016. DADA2: High-resolution sample inference from Illumina amplicon data. Nat Meth 13:581–583. https://doi.org/10.1038/nmeth.3869

52. Bokulich NA, Kaehler BD, Rideout JR, Dillon M, Bolyen E, Knight R, Huttley GA, Caporaso JG. 2018. Optimizing taxonomic classification of marker-gene amplicon sequences with QIIME 2’s q2-feature-classifier plugin. Microbiome 6: 90. https://doi.org/10.1186/s40168-018-0470-z

53. Robeson MS II, O’Rourke DR, Kaehler BD, Ziemski M, Dillon MR, Foster JT, Bokulich NA. RESCRIPt: Reproducible sequence taxonomy reference database management for the masses. bioRxiv 2020.10.05.326504. https://doi.org/10.1101/2020.10.05.326504

54. Edgar RC. 2004. MUSCLE: multiple sequence alignment with high accuracy and high throughput. Nucl Acids Res 32: 1792–1797. https://doi.org/10.1093/nar/gkh340

55. Ronquist F, Teslenko M, van der Mark P, Ayres DL, Darling A, Höhna S, Larget B, Liu L, Suchard MA, Huelsenbeck JP. 2012. MRBAYES 3.2: Efficient Bayesian phylogenetic inference and model selection across a large model space. Syst. Biol. 61:539–542. https://doi.org/10.1093/sysbio/sys029

